# MTFR2 regulates mitochondrial fission and impacts spindle integrity during mitosis

**DOI:** 10.1101/2020.09.13.293621

**Authors:** Yibo Luo, Song-Tao Liu

## Abstract

Previously we reported that mitochondrial fission regulator 2 (MTFR2, also termed DUFD1 or FAM54A) is co-transcribed with core centromere/kinetochore components, indicating a possible role in mitosis regulation. Here we show that human MTFR2 is a mitochondrial outer membrane protein and participates in DRP1 dependent mitochondrial fission. Multiple *MTFR2* variants identified in cancer samples are defective in triggering mitochondrial fission. Inducible *MTFR2* depletion caused prolonged mitotic duration and increased chromosome mis-segregation, resulting in multi-nucleated daughter cells. *MTFR2* knockout cells accumulated spindle defects, producing either multipolar spindles or short oscillating spindles due to loss of astral microtubules. MTFR2 is phosphorylated during mitosis. The phosphorylation mutant, as well as the cancer variants, failed to correct the prolonged mitotic duration. *MTFR2* knockout also rendered cells more resistant to apoptosis caused by taxol treatment. As overexpressing MFN1 or DRP1-K38A also caused spindle defects, we conclude that mitochondrial fragmentation during mitosis ensures spindle integrity and chromosomal stability, and MTFR2 plays a critical role in bridging proper mitochondrial fission and chromosome segregation.

## Introduction

Mitosis is more than segregating duplicated sister chromatids into daughter cells. Intracellular organelles also need to be properly distributed in order to produce healthy progenies. There is growing interest in understanding the crosstalk and coordination between chromosome segregation and inheritance of membrane organelles during mitosis (Colanzi and Corda 2007, Wang and Seemann 2011, Mishra and Chan 2014, Jongsma, Berlin et al. 2015, Guizzunti and Seemann 2016, Asare, Levorse et al. 2017, Gruneberg and Barr 2017, Champion, Pawar et al. 2019, Carlton, Jones et al. 2020). Compared to other organelles, mitochondria carry a small but significant fraction of eukaryotic genomes, making their inheritance during mitosis even more important. As a hub for metabolism, energy production and apoptosis, mitochondria also play important roles in integrating intrinsic and external signals to determine mitotic progression rates and cell fates (Perfettini, Roumier et al. 2005, Youle and Karbowski 2005, McBride, Neuspiel et al. 2006, Westermann 2010, Mishra and Chan 2014, Vyas, Zaganjor et al. 2016, Ruan, Lim et al. 2018).

In animal cells, mitochondrial fission activity is increased during mitosis to facilitate fragmentation and distribution of mitochondria into daughter cells. The increase is caused by mitotic phosphorylation of several known mitochondrial fission regulators such as DRP1, RALA and RALBP1 (Taguchi, Ishihara et al. 2007, Kashatus and Counter 2011, Kashatus, Lim et al. 2011). The dynamin-like GTPase DRP1 is the primary regulator of mitochondrial fission (Smirnova, Shurland et al. 1998, Bleazard, McCaffery et al. 1999, Ingerman, Perkins et al. 2005). DRP1 mostly resides in the cytoplasm but is recruited to mitochondria by MFF and MID49/MID51 to form contractile rings that constrict for mitochondrial fission (Otera, Wang et al. 2010, Loson, Song et al. 2013). However, mitochondrial fission is a dynamic multi-step process and involves many other proteins (Otera, Ishihara et al. 2013). How these additional proteins work with DRP1 especially during mitosis is largely unknown. Furthermore, despite some prior knowledge whether mitochondrial fission has direct impact on chromosome segregation as Golgi apparatuses does in mitosis also remains to be further characterized (Colanzi and Corda 2007, Mitra, Wunder et al. 2009, Qian, Choi et al. 2012, Guizzunti and Seemann 2016).

Mitochondrial Fission Regulator 2 (MTFR2, also named FAM54A or DUFD1) was retrieved when we searched for genes co-transcribed with a set of 64 centromere/kinetochore components (Tipton, Wang et al. 2012). The co-expression with well-characterized mitosis regulators was not detected with other mitochondrial fission/fusion regulators. Previously, mouse MTFR2 was shown to localize at mitochondria and affect mitochondrial morphology but the working mechanisms were not investigated (Monticone, Panfoli et al. 2010). In this report, we demonstrated that MTFR2 regulates mitochondrial fission in a DRP1-dependent manner. We also found that many cancer-occurring MTFR2 variants are loss-of-function in nature. Surprisingly, MTFR2 knockout by inducible Cas9 mediated CRISPR resulted in centrosome defects and loss of astral microtubules during mitosis, causing spindle instability and chromosome mis-segregation. Our results suggest that MTFR2 coordinates mitochondrial fission and chromosome segregation during mitosis.

## Results

### MTFR2 is co-expressed with mitosis regulators

We have previously searched for potential novel mitosis regulators by examining genes co-transcribed with a group of 64 centromere/kinetochore proteins (Tipton, Wang et al. 2012). MTFR2, as a mitochondrial protein, was surprisingly retrieved in the search based on transcriptional profiling datasets housed in both Gene Sorter and COXPRESdb (Kent, Hsu et al. 2005, Okamura, Aoki et al. 2015). Conversely, pathway analysis of the genes co-expressed with *MTFR2* identified “Cell Cycle” as the most enriched biological process, and such a pattern is conserved among human, mouse, dog, chicken and zebrafish. More interestingly, the co-expression with mitosis genes is not observed with other well-known mitochondrial fission/fusion genes including *FIS1, DNM1L* (encoding DRP1), *MFF, MIEF2 (*encoding MID49*), MIEF1* (encoding MID51), *RALA, RALBP1, MFN1, MFN2 and OPA1*. MTFR2 has been reported to associate with chromosomal instability (CIN) and its abnormal expression or mutants have been observed in cancer samples (Knosel, Schluns et al. 2003, Staub, Grone et al. 2006, Turashvili, Bouchal et al. 2007, Cheng, Ou Yang et al. 2013, Wang, Xie et al. 2018, Lu, Lai et al. 2019). Taken together, MTFR2 was associated with mitosis and cancer development, but how it works in these processes is unclear.

### MTFR2 is a mitochondrial outer membrane protein and promotes mitochondrial fission

Since little is known about MTFR2 (Monticone, Panfoli et al. 2010), we started by examining the subcellular localization of human MTFR2. Immunofluorescence in HeLa cells showed that MTFR2 mostly decorated mitochondria marked by Mito-tracker Red (Pearson’s correlation coefficient r=0.849) (Supplemental fig.1A). Similarly, a C-terminal GFP fusion of MTFR2 (MTFR2-GFP) surrounded Mito-tracker signals in both interphase and mitotic cells (Supplemental fig.1B&1C). Western blots of cell fractionation further confirmed that MTFR2 is primarily localized in the mitochondria (Fig.1A) and is an integral membrane protein since MTFR2 could be extracted by 1% Triton X-100 but not by 1M Na_2_CO_3_ (pH=11.5) (Fig.1B). In addition, MTFR2 was as sensitive to proteinase K treatment as mitochondrial outer membrane protein TOM20 but not as inner membrane protein TIM23 (Fig.1C). Therefore, our results indicate that MTFR2 is a mitochondrial outer membrane protein.

It was suggested that MTFR2 participates in mitochondrial fission (Monticone, Panfoli et al. 2010). To investigate this further, we expressed MTFR2-GFP in HeLa cells that were synchronized at the G1/S boundary by thymidine treatment. G1/S cells usually have elongated tubular mitochondrial network (Mitra, Wunder et al. 2009) (Fig.1D, untransfected). Mitochondrial morphology in MTFR2-GFP transfected cells was classified into three categories based on the average mitochondrial length (Supplemental fig. 1D&1E). Approximately 70.0±5.9% (n=40) of MTFR2-GFP positive cells showed mitochondrial hyper-fission, and 27.8±5.4% cells showed apparent mitochondrial fission, in contrast to the 0 and 7.3±2.5% in untransfected cells (Fig.1D middle row, Fig 1E). When MTFR2-GFP and mCherry-TOM20 expressing cells were fused by PEG treatment, fused cells exhibiting mitochondrial hyper-fission showed both GFP and mCherry signals in their mitochondria (Supplemental fig.2). This suggested that MTFR2 does not block mitochondrial fusion. However, the mitochondrial fragmentation caused by MTFR2-GFP expression was abolished when the cells were co-transfected with a dominant negative *DRP1-K38A* construct (Fig. 1D bottom row, Fig. 1E). The mutant renders DRP1 defective in its GTPase activity which is required for mitochondrial fission (Smirnova, Shurland et al. 1998, Smirnova, Griparic et al. 2001). The mitochondria in some co-transfected cells formed perinuclear clusters or “balloons” characteristic of the DRP1 mutant. These results supported that MTFR2 promotes DRP1-dependent mitochondrial fission.

**Figure 1:**
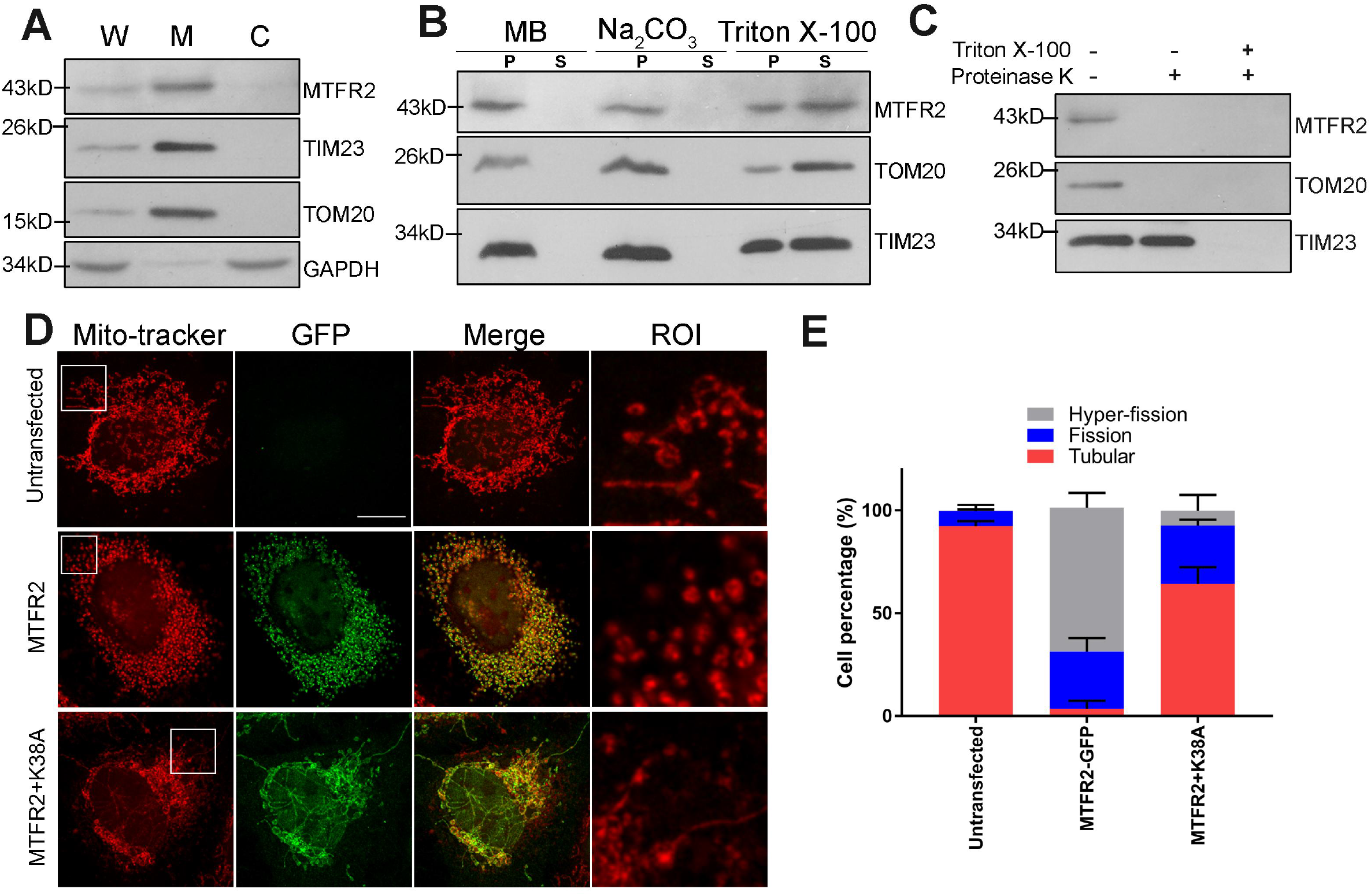
MTFR2 is a mitochondrial protein that promotes mitochondrial fission. (A) 50 µg of whole cell lysate (W), mitochondrial (M) and cytosolic (C) fractions were prepared from asynchronous HeLa cells and subjected to immunoblot. MTFR2 was probed together with mitochondrial inner membrane protein TIM23, mitochondrial outer membrane protein TOM20, and loading control GAPDH. Molecular weight markers (kD) were labelled on the left. (B) The mitochondrial fractions were re-suspended in mitochondrial buffer (MB) alone, or in MB plus 1M Na_2_CO_3_ (Ph=11.5) or 1% Triton X-100 in ice for 30 min. The insoluble membrane fragments (pellet, P) and supernatants (S) were probed for MTFR2, TIM23 and TOM20. (C) The mitochondrial fraction was re-suspended in MB and exposed to proteinase K. MTFR2 and outer membrane protein TOM20 were digested. Inner membrane protein TIM23 was left intact unless Triton X-100 was added. (D) MTFR2-GFP was transfected into Hela cells with or without *DRP1*-K38A co-expression (middle and lower rows) and the cells were synchronized at G1/S stage by 24hrs treatment of 2 Mm thymidine. Cells were stained with Mito-Tracker Red CMXRos for 30 min before fixation and imaged on Leica SP8 confocal microscope. Boxed areas were magnified for details. Bar=5µm. (E) Percentage of cells showing tubular, fission and hyper-fission mitochondrial morphology in untransfected, MTFR2-GFP or (MTFR2-GFP + DRP1-K38A) transfected HeLa cells (n=40, mean±SEM were from three independent experiments.).

### MTFR2 cancer-occurring variants are defective in inducing mitochondrial fission

To better understand the localization and activity of MTFR2, we mapped the mitochondrial targeting sequence of MTFR2 to its N-terminal (1-50) residues (Fig. 2A, Supplemental fig. 3A). It was predicted that an alternative spliced isoform b of MTFR2 missed the N-terminal 43 residues. This isoform of MTFR2 could not localize to mitochondria but enriched in the nucleus (Supplemental fig. 3A, third row), probably due to a stretch of basic residues at the C-terminus (299-302 residues, “KRKR”) that is usually masked by mitochondria targeting. We did not study this MTFR2 isoform further but attempted to map the MTFR2 region that is essential for the mitochondrial fission activity. MTFR2 contains a poly-proline sequence (PPS, 208-216 residues), which is conserved in its paralog MTFR1 and critical for the mitochondrial fission activity of MTFR1 (Tonachini, Monticone et al. 2004) (Fig. 2A). An MTFR2 mutant without this poly-proline sequence could not induce mitochondrial hyper-fission when overexpressed. Additional MTFR2 truncations fused with the mitochondrial targeting sequence (1-50), including (126-385), (126-343) and (126-290), or other constructs could not induce mitochondrial fragmentation as well (Supplemental fig.3B&C). We conclude that the fission activity of MTFR2 requires the intact protein or requires motifs scattered along the linear sequences.

**Figure 2:**
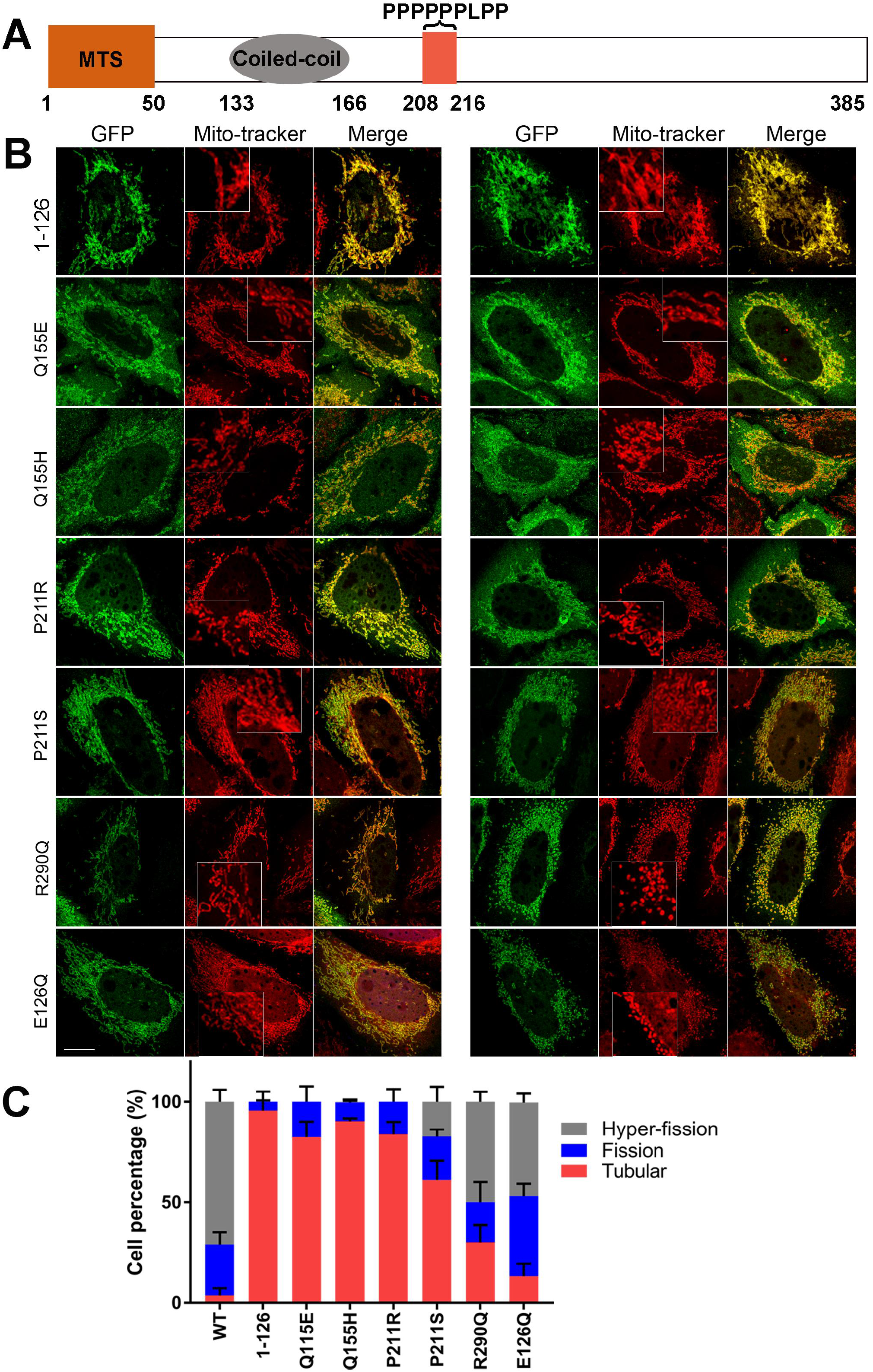
MTFR2 cancer-occurring variants are defective in mitochondrial fission. (A) Diagrammatic structure of MTFR2. The mitochondria targeting sequence (MTS, 1-50 residues), a predicted coiled-coil (133-166 residues) and the poly-proline sequence (208-216) are shown. (B) MTFR2-GFP harboring cancer-occurring variants were transfected into HeLa cells. Portion of the figure was enlarged to demonstrate the different mitochondrial morphology in MTFR2 variants. These cells were synchronized at the G1/S boundary for 24hrs after transfection. Bar=5μm. (C) Percentage of cells showing the fission activity after expression of wild type MTFR2-GFP or cancer-occurring variants (n=40, mean±SEM were from three independent experiments.)

**Figure 3:**
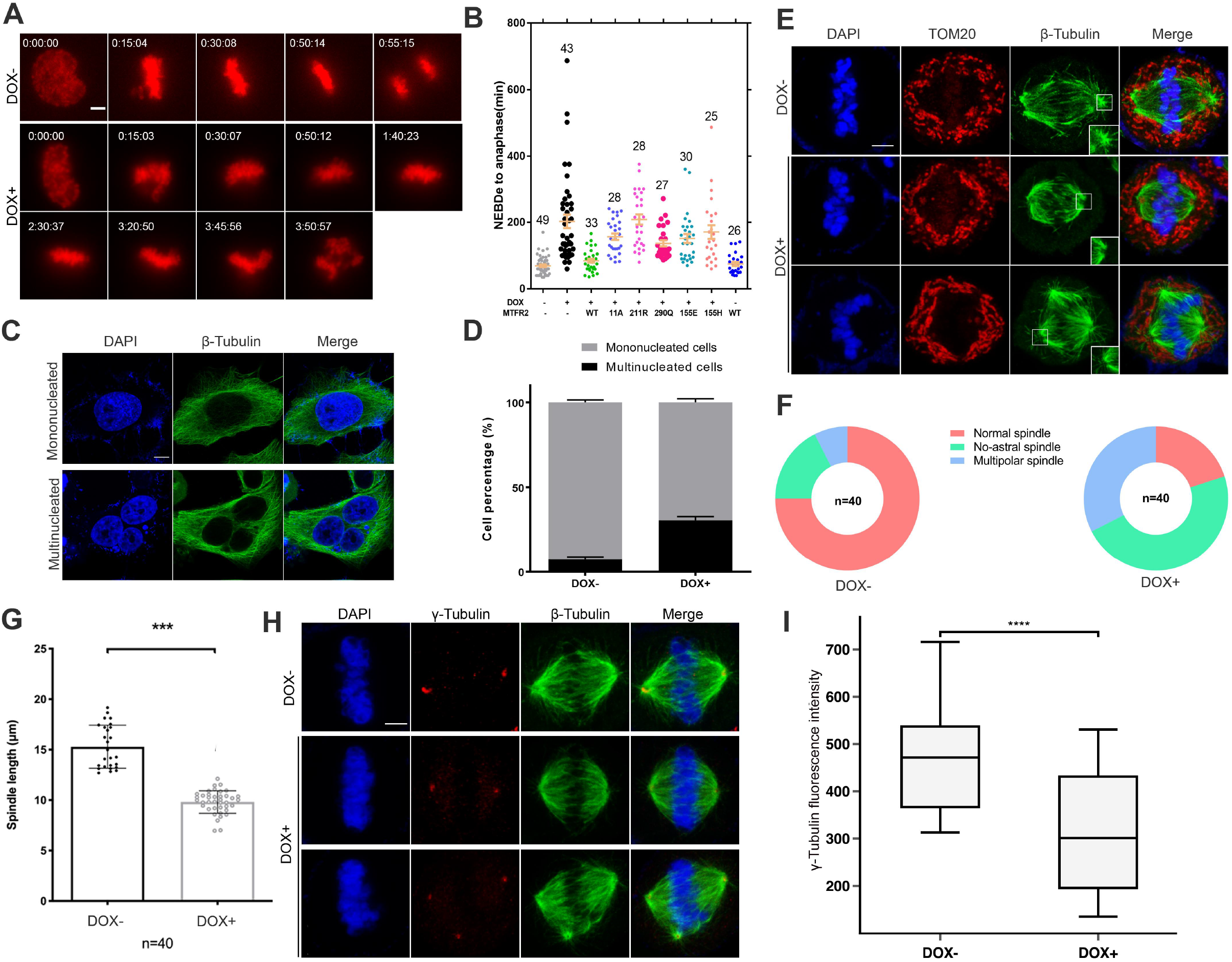
MTFR2 depletion results in mitotic spindle defects and chromosome mis-segregation. (A) Doxycycline was added for 48hrs to induce MTFR2 knockout in HeLa cells that inducibly express Cas9. Live cell imaging was conducted to compare mitotic durations in control (DOX-) and MTFR2 knockout (DOX+) cells that express Mrfp-histone H2A. Bar=5μm. (B) Mitotic durations were shown for *MTFR2* knockout that had been transfected with different MTFR2 constructs. The mitotic duration is defined as the time from nuclear envelope breakdown (NEBD) to anaphase onset. (C) Cells were released from RO3306 arrest. After 3h when most cells entered G1 stage, cells were fixed and strained with β-Tubulin and DAPI and imaged. Multinucleated cells were frequently found in MTFR2 knockout cells. Bar=5μm. (D) Quantification of multinucleated G1 cells after mitosis in control and *MTFR2* knockout cells. (n=40, mean± SEM was shown based on three independent experiments.). (E) Immunofluorescence of control (DOX-) and *MTFR2* knockout (DOX+) cells that were arrested in metaphase by treatment with MG132. Mitochondria were stained with anti-TOM20 antibody, while the spindles were stained with anti-β-Tubulin antibody. DNA was counterstained by DAPI. Enlarged portions compared the spindle poles and astral microtubules in the control and *MTFR2* knockout cells. Bar=5μm. (F) Pie graph indicating the percentage of mitotic cells showing multipolar spindles, spindles with no astral microtubules, or relatively normal bipolar spindles in control and *MTFR2* knockout cells (three independent experiments combined). (G) Quantification of the pole to pole spindle length of bipolar spindles in control and *MTFR2* knockout cells. *** indicates P<0.001 (Student’s t-test). (H) Immunofluorescence of control and *MTFR2* knockout cells stained with anti-γ-Tubulin and anti-β-Tubulin antibodies. (I) Quantification of γ-Tubulin fluorescence intensity at spindle poles of control and *MTFR2* knockout cells (cell number=24).

We wondered whether any MTFR2 point mutations could shed more light on the important portion of the protein for mitochondrial fission. We retrieved MTFR2 sequence variants in cancer samples from cBIOPORTAL and TumorPortal (Cerami, Gao et al. 2012, Gao, Aksoy et al. 2013, Lawrence, Stojanov et al. 2014). We chose to focus on (a) MTFR2 truncations caused by nonsense or splicing mutations, or (b) recurring MTFR2 mutants at the same residue in multiple samples and having at least “medium” impact judged by Mutation Assessor (Reva, Antipin et al. 2011) (Supplemental Table 1). Results showed none of these mutations could induce mitochondrial fragmentation as wild type did, except that E126Q found in renal clear cell carcinoma and R290Q found in colorectal cancer had over 50% of transfected cells showing fragmented mitochondria (“fission” and “hyper-fission” phenotype) (Fig.2B&2C, Supplemental fig.3C). Interestingly, Q155 and P211 fall in a predicted coiled coil (133-166) and the poly-proline sequence motifs respectively. These experiments suggested that cancer-occurring MTFR2 variants tend to be loss-of-function mutants.

### MTFR2 is not required for DRP1 targeting to mitochondria

To further test whether MTFR2 is required for mitochondrial fission, we established several HeLa cell-lines that depleted MTFR2 by CRISPR with inducible Cas9. After 48hrs treatment with doxycycline, MTFR2 was undetectable by western blot (Supplemental fig.4A). *MTFR2* knockout cells and control cells (without doxycycline) were arrested at metaphase with MG132. In control cells, mitochondria were fragmented into puncta as reported before (Taguchi, Ishihara et al. 2007, Kashatus, Lim et al. 2011), while in *MTFR2* knockout cells mitochondria were still forming the tubular network morphology, as shown by the longer mitochondrial length than in control cells (Supplemental fig.4B&C). In addition, the junctions connecting proximity mitochondrial was increased from 12 in control cells to 291 in knockout cells, suggesting abnormality of mitochondrial fission after MTFR2 knockout. However, in interphase cells, there was no significant difference in either mitochondrial length or network junction numbers between control and *MTFR2* knockout cells (Supplemental fig.4D&E). The results suggested basal level of MTFR2 is more critical in maintaining mitochondrial fragmentation during mitosis than in interphase cells.

Several proteins such as MFF, MID49 and MID51 facilitate mitochondrial fission by recruiting DRP1 from the cytoplasm to the mitochondria (Otera, Wang et al. 2010, Palmer, Osellame et al. 2011, Zhao, Liu et al. 2011, Loson, Song et al. 2013). The overall DRP1 level was not altered in *MTFR2* knockout cells (Supplemental fig.4A). To test whether MTFR2 is required for DRP1 recruitment to mitochondria, we measured the DRP1 signals overlapping with mitochondria. The overlapping signals were higher in *MTFR2* knockout cells than in control cells (Supplemental fig.5A&B). The result was confirmed by western blot showing more DRP1 in isolated mitochondrial fraction from *MTFR2* knockout cells (Supplemental fig.5C). In addition, direct interaction of DRP1 and MTFR2 was barely detected by GFP-Trap (Supplemental fig.5D). The results indicate that MTFR2 is not required for recruiting DRP1 to mitochondria despite that its mitochondrial fission activity depends on DRP1 (Fig.1D).

### MTFR2 is phosphorylated during mitosis

We next investigated whether MTFR2 protein undergoes any modifications during mitosis since MTFR2 is co-transcribed with core centromere/kinetochore proteins (Tipton, Wang et al. 2012). The cell lysates were prepared from HeLa cells synchronized at G1/S, G2/M and prometaphase by treatment with thymidine, RO3306 and nocodazole, respectively. Comparing the immunoblots of the lysates, there was accumulation of MTFR2 protein during mitosis, which also exhibited a dramatic mobility shift (Supplemental fig.6A). Lambda-Phosphatase treatment further showed that the mobility shift was caused by phosphorylation (Supplemental fig.6B). Samples taken from cells released from nocodazole demonstrated that the MTFR2 mobility shift diminished along with the degradation of CYCLIN B1 (Supplemental fig.6C), also indicating that MTFR2 phosphorylation is mitosis specific. Next, we tried to detect which mitotic kinases are responsible for MTFR2 phosphorylation by applying small molecule inhibitors to cells already arrested in mitosis with nocodazole and the proteasome inhibitor MG132 (to prevent mitotic slippage). All kinase inhibitors reduced MTFR2 mobility shift except the MPS1 inhibitor reversine, even at high concentrations (Supplemental fig.7A, B&C). We noticed that residual MTFR2 mobility shift was reproducibly detected if PLK1 inhibitor BI2536 was applied onto nocodazole arrested prometaphase cells (Supplemental fig.7D), but the shift disappeared completely if BI2536 was added directly into thymidine released cells (Supplemental fig.7E). The above results suggest that mitotic phosphorylation of MTFR2 is affected directly and indirectly by CDK1, AURORA and PLK1 kinases but not MPS1 kinase.

### MTFR2 depletion leads to mitotic defects

Live cell imaging was performed on control and MTFR2 knockout cells expressing mRFP-histone H2A. The mitotic duration from nuclear envelope breakdown to anaphase onset in control cells was 69.3±3.9 min (Fig 3A&B). However, MTFR2 depletion significantly increased this duration to 202.8±20.1 min (P<0.001, Student’s t-test) (Fig.3A&3B, Supplemental videos 1 and 2). The chromosome mis-segregation increased in the MTFR2 knockout cells, giving rise to 30.6% multi-nucleated cells following cell divisions, significantly more than 7.4% in control cells (Fig.3C&3D).

The prolonged mitotic durations in MTFR2 knockout cells were corrected by rescuing with expression of wild type MTFR2 fused with GFP (WT) (Fig. 3B). Cancer-occurring MTFR2 variants Q155E, Q155H, P211R, and R290Q could not rescue the mitotic duration defect. In addition, we searched PhosphositePlus for in vivo phosphorylation sites of MTFR2 (Hornbeck, Zhang et al. 2015), and generated an 11A mutant, which mutated 11 serine/threonine residues into alanine (Material and Methods; Supplemental fig. 1C,11A). The 11A mutant could not rescue the mitotic duration defect either, in which the mitotic duration from NEBD to anaphase was 146.4±48.3min (Fig.3B).

We went on to examine the underlying reasons of the mitotic delay. There were no apparent differences in control and *MTFR2* knockout cells regarding the cold-stable kinetochore microtubules nor the frequency of chromosome misalignment in MG132 arrested metaphase (data not shown), indicating that the kinetochore-microtubule attachment was not grossly affected by *MTFR2* knockout. However, live cell imaging of mCherry-Tubulin transfected cells detected that spindle oscillation was frequent in *MTFR2* knockout cells, although the spindle was usually at the cell center when chromosomes started to segregate (Supplemental fig.8A&B, Supplemental videos 3 and 4). Immunofluorescence confirmed two major spindle defects in *MTFR2* knockout mitotic cells: around 32.5% of cells harboring multi-polar spindles, and 47.5% lacking astral microtubules (Fig.3 E&F), in contrast to 7.5% and 20.0% in control cells. The cells lacking astral microtubules tend to have shorter spindles (Fig. 3G). MTFR2 knockout cells also showed lower γ-Tubulin intensity at spindle poles (Fig. 3H&I), indicating reduced microtubule nucleating activities. Some spindle poles did not have any or only have one CENTRIN-1 dot associating with γ-Tubulin foci in MTFR2 knockout cells (Supplemental fig. 8C&D), indicating centriole splitting or loss in these cells. Collectively, the above results demonstrated that MTFR2 knockout caused a range of spindle defects that resulted in chromosome mis-segregation.

### Blocking mitotic mitochondrial fragmentation results in spindle defects

Previous reports also suggested that chromosome mis-segregation and centrosome amplification could happen after *DRP1* knockdown, without detailed characterization of the mitotic spindle (Mitra, Wunder et al. 2009,Qian, Choi et al. 2012). To further test the potential connections between mitotic mitochondrial fragmentation and the spindle integrity, we transfected mitochondrial fusion factor *MFN1* or the dominant negative *DRP1-*K38A mutant along with mCherry-*TOM20* into Hela cells and checked the spindle formation in MG132 arrested cells.

Similar as in *MTFR2* knockout cells, *MFN1* or *DRP1*-K38A transfection induced longer tubular mitochondrial network during mitosis (Fig.4A&B). Interestingly, 53.3% and 33.3% of cells showed multi-polar spindles in *MFN1* and *DRP1*-K38A transfected cells, respectively, compared 7.5% and 32.5% in control and MTFR2 knockout cells (Fig. 3F&4C). These results support that mitochondrial fission defects impact mitotic spindle integrity.

### MTFR2 knockout renders HeLa cells taxol resistant

Ablation of mitochondrial fission factors usually but not always renders affected cells apoptosis resistant (Lee, Jeong et al. 2004, Perfettini, Roumier et al. 2005). We subjected MTFR2 knockout cells to taxol treatment and the imaging results suggested the knockout cells were indeed resistant to taxol triggered apoptosis (Fig.4D). Both control and MTFR2 knockout cells were released from thymidine block into taxol-containing medium. Cells were imaged 8 hrs later when they started entering mitosis (see Methods). Some control cells underwent membrane blebbing and cell death as early as 7 hrs into the imaging. However, in *MTFR2* knockout cells, 50% of the cells were still alive after 24hrs (Fig. 4E and Supplemental movie 5&6). This result shows that MTFR2 knockout renders HeLa cells resistant to taxol-induced apoptosis. Interestingly, preliminary tests found both control and knockout cells were still sensitive to doxorubicin, a DNA topoisomerase II inhibitor, induced apoptosis, suggesting mitotic specific function of MTFR2 (data not shown).

**Figure 4:**
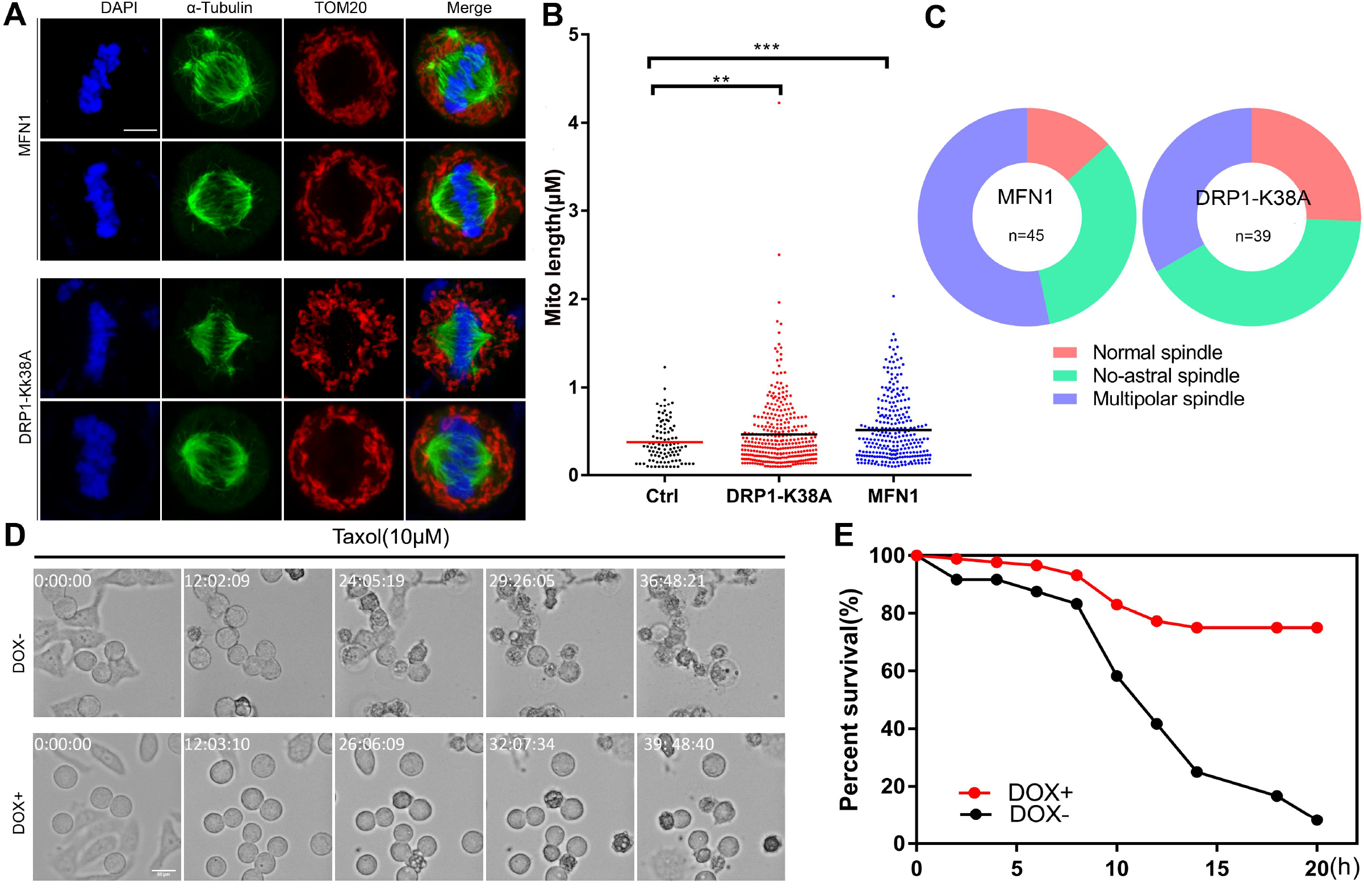
Mitotic mitochondrial fragmentation affects the integrity of mitotic spindles and taxol resistance. (A) HeLa cells were co-transfected with *MFN1* or *DRP1*-K38A and mCherry-*TOM20* (3:1), arrested in mitosis by MG132 treatment and stained for β-Tubulin. Shown were the spindles lacking astral microtubules or multi-polar spindles. Bar=5μm. (B) The mitochondrial length in control (Ctrl), DRP1-K38A and MFN1 transfected cells. (C) Pie graph showing the percentage of cells with no astral microtubules, with multipolar spindles or with relatively normal spindles (three independent experiments combined.) (D) Doxycycline and thymidine were added at the same time. After 24hrs, cells were released into fresh medium containing 10µM taxol. Live cell imaging started after 8hrs when cells were entering mitosis (marked as time 0). Images for control and MTFR2 knockout cells at different time points were shown. The time stamp is in hr:min:sec after imaging. (E) Quantitation of percent (%) cell survival (n=100) based on the movies shown in (Supplemental movie 5&6).

## Discussion

We showed that MTFR2 mitochondrial fission activity depends on DRP1 (Fig. 1), but MTFR2 does not function as a mitochondrial receptor for DRP1 like MFF, MID51 or MID49 (Supplemental fig. 5) (Otera, Wang et al. 2010, Palmer, Osellame et al. 2011, Zhao, Liu et al. 2011, Loson, Song et al. 2013). As the essential GTPase for mitochondrial constriction and eventual scission, DRP1 activity is regulated by its mitochondrial recruitment, polymerization status, interactions with accessory proteins and lipid components, with many of the interactions transient (Tandler, Hoppel et al. 2018). Importantly, DRP1 polymerization usually stimulates its GTPase activity; however, previous work clearly showed that DRP1 can form different types of oligomers and not all of them are conducive for mitochondrial fission (Mears, Lackner et al. 2011, Francy, Alvarez et al. 2015, Clinton, Francy et al. 2016, Francy, Clinton et al. 2017, Kalia, Wang et al. 2018, Hoppins, Lackner et al. 2020). Future work will test whether MTFR2 especially the phosphorylated MTFR2 during mitosis helps to guide the formation of productive DRP1 oligomers on mitochondria or enhance DRP1 GTPase activity to increase mitochondrial fission.

The mitotic spindle defects we observed when mitotic mitochondrial fragmentation was compromised (Fig. 3&Fig. 4) extended previous reports with *DRP1* knockdown (Mitra, Wunder et al. 2009, Qian, Choi et al. 2012). It should be noted that defects in Golgi and peroxisome organization and localization in mitotic cells sent retrograde signaling to affect the orientation and stability of the mitotic spindle (Colanzi and Corda 2007, Guizzunti and Seemann 2016, Asare, Levorse et al. 2017). Therefore, the evidence is accumulating to support the coordination between chromosome segregation and the inheritance of membrane organelles (Wang and Seemann 2011, Jongsma, Berlin et al. 2015, Carlton, Jones et al. 2020). There were suggestions that mitochondrial hyper-fusion as caused by DRP1 knockdown increases cyclin E level and accumulates DNA damage that eventually impacted chromosome segregation in subsequent mitosis (Mitra, Wunder et al. 2009, Qian, Choi et al. 2012). Our results here strongly supported that the integrity of mitotic spindles was often compromised when mitochondrial fragmentation failed to be executed during mitosis. Whether the loss of astral microtubules and the appearance of multipolar spindles are inherently connected and what are the underlying molecular mechanisms are topics for future investigation.

Imbalance of mitochondrial fission and fusion has been observed during oncogenesis and cancer development, although the exact pathological impact may be context-dependent (Vyas, Zaganjor et al. 2016). We found that multiple cancer occurring MTFR2 variants both compromised mitochondrial fission and failed to rescue mitotic defects (Fig 2, 3). This raised the possibility that these sporadic MTFR2 variants facilitate cancer progression by increasing chromosomal instability. In addition, as MTFR2 depletion enhanced taxol resistance, the cells without fully functional MTFR2 could survive at least some of the cancer drugs. The long-term benefits of these MTFR2 variants in cancer cell proliferation, evolution and survival will be addressed in follow-up studies.

In conclusion, we have showed that MTFR2 promotes mitochondrial fission particularly during mitosis. Interfering with mitotic mitochondrial fragmentation by MTFR2 knockout or *MFN1* or *DRP1*-K38A overexpression accumulates mitotic spindle defects, delaying mitotic progression and increasing chromosome mis-segregation. MTFR2 loss also enhances taxol resistance. We suggest that MTFR2 plays a crucial role in coordinating faithful chromosome segregation with mitochondrial inheritance and its disruption benefits cancer cells.

## Material and methods

### 1 Cell culture, knockout cell line generation and drugs

Hela cells were maintained in DMEM (Corning) supplemented with 10% FBS (Corning), L-glutamine and 1X non-essential amino acids (GE) and cultured under 5% CO_2_ at 37^°^C (Ji, Luo et al. 2018). The HeLa cell line with doxycycline inducible Cas9 (cTT20.11) was kindly provided by Dr. Iain Cheeseman (MIT) (McKinley and Cheeseman 2017).

To prepare single guide RNA targeting MTFR2, DNA oligoes (F: CACCGACGACATTTACCTGTTCTAC, R: AAACGTAGAACAGGTAAATGTCGTC) were annealed and cloned into BsmBI (NEB) cut pLenti-sgRNA to make pLenti-sgMTFR2 following published protocols (Ran, Hsu et al. 2013). The pLenti-sgMTFR2 plasmid as well as lentivirus packaging plasmid psPAX2 and envelope plasmid pMD2.G (ratio was 10:9:1) were co-transfected into 293-FT cells, with medium change once after 24hrs. After 48hrs, the virus was collected by harvesting and filtering the medium. The lentivirus-containing filtrate along with 8μg/ml polybrene was added into 50% confluent cTT20.11 cells, which had been cultured for at least 24hrs. Single cell colonies were isolated and expanded after selection with puromycin (Su, Tsang et al. 2018). To knockout MTFR2, cells were seeded in the culture medium containing 1µg/ml doxycycline for 48 hr.

To arrest cells at different cell cycle stages, 2 mM thymidine (Sigma) was applied for 24hrs to arrest cells at G1/S boundary; 10 µM RO3306 (EMD millipore) treatment for at least 6hrs was used to arrest cells at G2/M boundary; 330 nM nocodazole (Sigma) was employed to arrest cells at prometaphase; and 10µM MG132 was used to inhibit anaphase onset (Ji, Arnst et al. 2016).

PEG mediated cell fusion was done by previously reported (Zhao, Liu et al. 2011). Briefly, MTFR2-GFP or mCherry-TOM20 transfected cells were trypsinized, washed with PBS and mixed. The mixed cells were treated with 50% PEG for 10min under room temperature. Then the cells were washed with PBS again and seeded on a new plate. 24hrs after fusion, the cells were fixed and subjected on confocal to detect the GFP and mCherry double positive cells.

### 2 PCR and DNA constructs

The full length *MTFR2* and various truncations were PCR amplified from a HeLa cell cDNA preparation, and subcloned in pENTR using the pENTR/D-TOPO cloning kit (Thermofisher Scientific). Site-direct mutagenesis of *MTFR2* was generated using a pair of complementary primer sets, and PCR was done with Q5 High Fidelity DNA polymerase (NEB). The LR reaction kit (Thermofisher Scientific) was used to subclone the *MTFR2* fragments into pDEST47, which fused GFP at the C-terminal of *MTFR2* or its truncations/mutants. In the MTFR2-11A phosphor-mutant, the 11 serine/threonine sites are S119, S291, S305, S311, S314, S328, S335, S342, S354, T378, and S379. mRFP-Histone H2A and mCherry-Tubulin plasmids were described before (Wang, Sturt-Gillespie et al. 2014). Other plasmids from Addgene were listed in Supplemental Table 2. All constructs were confirmed by DNA sequencing (Genewiz).

### 3 Cell transfection, immunofluorescence and live cell imaging

Polyethylenimine (PEI) was used in DNA transfection as described before (Ji, Arnst et al. 2016). Briefly, for a 3.5 cm dish, 0.5 µl PEI (5µg/µl) and 2µg DNA was mixed with 200 µl Opti-MEM, and the mixture was incubated at room temperature for 30 min before applying dropwise to cells.

For immunofluorescence, cells were grown on poly-L-lysine coated glass coverslips and fixed with 4% paraformaldehyde (PFA) for 10 min at room temperature (37^°^C for β-Tubulin staining). For γ-Tubulin and CENTRIN-1 staining, cells were fixed with 100% methanol at 37^°^C for 10 min. The fixed cells were extracted by KBT (buffer for permeabilization: 10 mM Tris at pH7.5, 150 mM NaCl, 0.1%BSA, 0.2% Triton X-100) for 2 min. After blocking in KB (buffer for blocking, KBT minus Triton X-100) for at least 5 min, primary antibodies were diluted in KB and incubated at 4^°^C overnight or 37^°^C for 1hr. Secondary antibodies conjugated with Alexa Fluor 488, 555 and 647 (Life technologies) were used at 1:1000 dilution in KB. DNA was stained with DAPI-containing Fluorosheild mounting medium (Sigma). Images were acquired on a Leica SP8 confocal microscope usually using 100 × objective (NA=1.40) as 1 µm Z-stacks. The primary antibodies used were listed in Supplemental Table 3. For mitochondrial labeling, mito-trakcer Red CMXRos (Thermofisher Scientific) was added into culture medium directly 30min before cell fixation. The final concentration is 500nM.

Live cell imaging for mitotic duration analyses for Fig. 4 was carried out on an Olympus IX81 epifluorescence microscope essentially as before (Wang, Sturt-Gillespie et al. 2014), except that images were acquired with a 60 × objective (NA=1.42) at intervals of 5 min. Live cell imaging for spindle oscillation for Supplemental fig. 8 was performed on Leica SP8 confocal microscope using 100 × objective at 1 µm Z-stacks with 3 min intervals.

### 4 Quantitative microscopy image analysis

Image J(Fiji) was employed for quantitative image analyses (Schindelin, Arganda-Carreras et al. 2012).To determine the mitochondrial length and network formation, default threshold was applied to each image and the images were then skeletonized by Plugin “Skeletonize (2D/3D)”. The “Ridge detection” plugin was used to quantify the number of junctions and the length between each junction or separated single mitochondria (Steger 1998). The “JACoP” plugin was used to quantify the fractions of DRP1 associated with mitochondria (Bolte and Cordelieres 2006). The same threshold was applied to all images and the Manders’ overlap coefficients were used to compare the DRP1 associated with mitochondria in different conditions. The Pearson’s correlation coefficients were used to quantify the MTFR2 co-localization with mitochondria.

### 5 Cell fractionation, GFP-Trap, lambda-phosphatase treatment and immunoblot

To isolate mitochondrial fraction, Hela cells were harvested in ice-cold mitochondrial buffer (MB, 210 mM mannitol, 70 mM sucrose, 10 mM HEPES, 1 mM EDTA, pH 7.5) supplemented with proteinase inhibitor (Roche). Cell suspension was stroked with a 27-gauge needle for 25 times and then was centrifuged at 1500g for 5min under 4^°^C. The resulting supernatant was re-centrifuged at 16,000g for 30min. The resulting supernatant is the cytosol fraction while the pellet is the mitochondrial fraction. For the membrane protein analysis, pellet mitochondrial fraction was resuspended in 100µl MB or MB containing 0.1M Na_2_CO_3_ (pH=11.5) or MB containing 1% Triton X-100 and incubated in ice for 30min. The insoluble membrane fractions were centrifuged at 16,000 g for 15 minutes, and the supernatants precipitated with 10% (v/v) trichloroacetic acid. For proteinase K treatment, pellet mitochondria were treated with 50µg/ml proteinase K in the absence or in the presence of 1% Triton-X 100 for 30min under room temperature. Centrifuging at 16,000g for 15min to pellet the insoluble membrane fraction.

Buffers for making cell lysates and immunoprecipitation were described as before (Tipton, Tipton et al. 2011). 300 µg of cell lysate was used for GFP-Trap with 20 µl beads (Chromotek) (50% slurry). The immunoprecipitants were washed and 10 µl 2X SDS-PAGE loading buffer was added. For lambda-phosphatase treatment, 50 µg cell lysate was treated with 80 units of lambda-phosphatase (NEB) at 30^°^C for 30 min, in the absence or presence of phosphatase inhibitors (mixture of 10mM NaF, 1mM Na_3_VO_4_ and 60mM β-glycerophosphate). The reactions were stopped by adding 1/3 volume of 4X SDS-PAGE loading buffer. The immunoblots were probed with proper primary antibodies, alkaline phosphatase-conjugated secondary antibodies (1:30,000 dilution, Thermofisher scientific), and Tropix CDP-star (Applied biosystems) as the luminescent substrate.

### 6 Statistical analysis

All experiments in this study were repeated at least three times. GraphPad Prism was used for statistical analyses. Comparison of control and experimental groups was usually analyzed by Student’s t-test, with P<0.05 considered as showing significant difference.

## Supplemental Data

8 Supplemental Figures

3 Supplemental Tables

6 Supplemental Videos

## Supplemental Figure Legends

*Supplemental figure 1: MTFR2 is a mitochondrial protein that promotes mitochondrial fission.*

(A) Immunofluorescence of an interphase HeLa cell. MTFR2 was detected by a rabbit anti-MTFR2 antibody and an Alexa Fluor 488 conjugated secondary antibody. Mitochondria were labelled by Mito-Tracker Red. DNA was counterstained with DAPI. A region of interest (ROI) marked by a square was magnified for details. Bar=5μm.

(B) *MTFR2*-GFP was transfected into HeLa cells. GFP and Mito-Tracker signals were shown in an interphase (top row) and a mitotic cell (bottom row). Bar=5μm.

(C) Immunoblot of 50 µg mitotic lysates from HeLa cells either untransfected or transfected with *MTFR2*-GFP or *MTFR2*-11A-GFP. MTFR2-GFP and endogenous MTFR2 were of expected sizes. MAD2 was probed as a loading control.

(D) Representative images of *MTFR2*-GFP-expressing cells to show three categories of mitochondrial morphologies: tubular, fission, and hyper-fission.

(E) Dot plots of the mitochondrial length in the three categories of cells as shown in (D). The mitochondrial length was determined as detailed in Materials and Methods. The average length is indicated by the short horizontal bar in each category. *** indicates P<0.01 (Student’s *t*-test).

*Supplemental figure 2: MTFR2-GFP expression does not interfere with mitochondrial fusion.*

HeLa cells were separately transfected with *MTFR2*-GFP and mCherry-*TOM20*. 6hrs after transfection, the cells were trypsinized, treated with PEG for cell fusion and re-plated in thymidine added culture medium. The cells were examined 24hrs later by immunofluorescence. Two fused cells with both GFP and mCherry signals in their mitochondria are shown in the lower portion, in which the mitochondria were also hyper-fission. Bar=5μm.

*Supplemental figure 3: Domain mapping of MTFR2.*

(A) The N-terminal 50 amino acids of MTFR2 is necessary and sufficient for mitochondria targeting. Note that (1-43), (44-385) and (10-50) residues of MTFR2 failed to target C-terminal fused GFP to mitochondria. The line plots on the right show whether GFP signals are coincident with Mito-tracker Red signals.

(B) Mitochondria were not fragmented when cells were transfected with MTFR2 missing the poly-proline sequence (PPS) or other MTFR2 truncations fused with the mitochondrial targeting sequence (1-50) (left side). Fragments expanding the first 60 or 80 amino acids as well as cancer mutations S172x and R290x expressing results are shown (right side).

(C) Summary of MTFR2 fragments and point mutations tested for mitochondrial localization and mitochondrial fission activity.

*Supplemental figure 4: MTFR2 knockout affects mitochondrial fission in mitotic cells.*

(A) HeLa cells that contain sgRNA targeting *MTFR2* and inducibly express Cas9 were treated with doxycycline (DOX) for 48hrs before harvesting for cell lysates. The control (DOX-) and knockout (DOX+) cell lysates were probed for MTFR2, DRP1 and TOM20.

(B) Immunofluorescence images to show mitochondrial morphology in control and *MTFR2* knockout metaphase cells arrested with MG132. Bar=5μm.

(C) Quantification of mitochondrial length in control and *MTFR2* knockout metaphase cells.

(D) Immunofluorescence images to show mitochondrial morphology in control and *MTFR2* knockout interphase cells. Cells were stained with anti-TOM20 and anti-β-Tubulin antibodies. Bar=5μm.

(E) Quantification of mitochondrial length in control and *MTFR2* knockout interphase cells.

*Supplemental figure 5: MTFR2 is not required for the mitochondrial recruitment of DRP1.*

(A) Immunofluorescence of control (DOX-) and *MTFR2* knockout (DOX+) metaphase cells stained with anti-TOM20 and anti-DRP1 antibodies. Zoomed areas (ROI) were shown for details of DRP1 associated with mitochondria. Bar=5μm.

(B) Quantification of DRP1 signals overlapped with mitochondria (defined by TOM20 signals) as in (A), expressed as Mander’s overlap coefficients.

(C) The whole cell lysates (WCL), mitochondria and cytosol fractions were prepared from control and *MTFR2* knockout mitotic cells and subjected to immunoblot. DRP1, MTFR2 as well as TIM23 and GAPDH were probed.

(D) Immunoblot followed by GFP-trap using cell lysates from control or MTFR2-GFP expressing cells. Direct interactions between MTFR2 and DRP1, PROFILIN, ACTIN was not detected.

*Supplemental figure 6: MTFR2 is phosphorylated during mitosis.*

(A) Immunoblot of MTFR2 in lysates from asynchronous HeLa cells or cells arrested at G1/S boundary, G2/M boundary and prometaphase by treatment with thymidine, RO3306 and nocodazole respectively. MAD2 was probed as a loading control.

(B) 50 µg lysates from interphase or mitotic HeLa cells were probed for MTFR2 and MAD2 by Western blot. The mitotic lysates were untreated or treated with λ-phosphatase in the absence (-) or presence (+) of phosphatase inhibitors. The numbers on the left indicate molecular weight markers (kD).

(C) Western blot of MTFR2, CYCLIN B1 and MAD2 of samples prepared at different time points after HeLa cells were released from nocodazole arrest.

*Supplemental figure 7: MTFR2 phosphorylation is regulated by mitotic kinases.*

(A, B, C, D) Increasing concentrations of Hesperadin (an inhibitor for AURORA A and B kinases), Reversine (MPS1 kinase inhibitor), RO3306 (a CDK1 inhibitor) and BI2536 (a PLK1 inhibitor) were added to cells already arrested in nocodazole and MG132 and incubated for 2hrs, and mitotic cells were harvested by shake-off and cell lysates were used for immunoblot of MTFR2. GAPDH were probed as loading controls.

(E) BI2536 was added into thymidine release cells for 12hrs. Cells were accumulated at mitosis because PLK1 inhibition blocks chromosome segregation. Mitotic cells were harvested by shake-off and cell lysate were subjected to immunoblot, which was probed as above.

*Supplemental figure 8: Mitotic defects in MTFR2 knockout cells.*

(A) HeLa cells engineered for inducible *MTFR2* knockout were transfected with mCherry-Tubulin, treated with doxycycline for 48hrs (DOX+) or not (DOX-), and imaged through mitosis. Time was marked as hr:min:sec. Bar=5μm.

(B) The distances from two spindle poles to the neighboring cell cortex were measured at each frame, and the ratio of the distances were plotted for three DOX- and three DOX+ cells.

(C) Immunofluorescence of control and *MTFR2* knockout cells with anti-γ-Tubulin and anti-CENTRIN-1 antibodies. Bar=5μm.

(D) Pie graph showing the quantification of CENTRIN numbers associated with each γ-Tubulin. While control cells exhibited two CENTRIN signal dots near one γ-Tubulin focus, in *MTFR2* knockout cells many γ-Tubulin foci associated with only one or none CENTRIN dot (three independent experiment combined.)

**Supplemental Table 1:** MTFR2 cancer-occurring variants tested in this work

| MTFR2 Mutations | Cancer Tissues |
| --- | --- |
| E126Q | Renal clear cell carcinoma |
| E126x | Uterine corpus endometrial carcinoma |
| Q155E | Head and neck cancer |
| Q155H | Liver cancer |
| S172x | Small cell lung cancer and acinar cell carcinoma of pancreas |
| P211R, P211S | Head and neck cancer |
| R290Q | Colorectal cancer |
| R290x | Lung adenocarcinoma and colorectal cancer |

*X: splicing truncations or nonsense mutations.*

**Supplemental Table 2:** Addgene plasmids

| Plasmid | Addgene ID | References |
| --- | --- | --- |
| mCherry-Drp1 | 49152 | (Friedman, Lackner et al. 2011) |
| mCherry-TOMM20-N-10 | 55146 | Michael Davidson Lab |
| Mfn1-Myc | 23212 | (Chen, Detmer et al. 2003) |
| pcDNA3-Drp1K38A | 45161 | (Smirnova, Shurland et al. 1998) |
| pLenti-sgRNA | 71409 | Eric Lander & David Sabatini |

---

**Supplemental Table 3:** Primary antibodies used in this study

| Target protein | Source | Use |
| --- | --- | --- |
| MTFR2 | Thermofisher scientific, PA5-56258, rabbit | IB: 1:1000<br>IF: 1:50 |
| DRP1 | BD Transduction Laboratories, 611738, mouse | IF: 1:200 |
| DRP1 | Abclonal, A2586, rabbit | IB: 1:500 |
| β-Tubulin | Cell signaling, 86298, mouse | IF: 1:200 |
| γ-Tubulin | Sigma-Aldrich, T5192, rabbit | IF: 1:50 |
| TOM20 | Abclonal, A6774, rabbit | IB: 1:1000<br>IF: 1:200 |
| TIM23 | Abclonal, A8688, rabbit | IB: 1:1000 |
| MAD2 | Bethyl A300-301A, rabbit, | IB: 1:500 |
| GFP | Invitrogen, A11122, rabbit | IF: 1:200 |
| mCherry | ThermoFisher Scientific, PA5- 34974, rabbit | IF: 1:100 |
| CENTRIN-1 | Santa Cruz biotechnology, SC-293494, mouse | IF: 1:15 |
| CYCLIN B | Santa cruz biotechnology, SC-245, mouse | IB: 1:500 |
| GAPDH | Abclonal, AC002, mouse | IB: 1:500 |

## Description of Supplemental Videos

*Supplemental video 1:*

The video shows a control HeLa cell transfected with mRFP-histone H2A going through mitosis. The nuclear envelope breakdown (NEBD) and separation of sister chromatids (anaphase onset) are monitored by chromatin dynamics as indicated by mRFP tagged histone H2A.

*Supplemental video 2:*

The video shows a *MTFR2* knockout HeLa cell transfected with mRFP-histone H2A going through mitosis.

*Supplemental video 3:*

The video shows a control HeLa cell transfected with mCherry-tubulin undergoing mitosis. The mCherry-tubulin signals are overlaid on the brightfield images.

*Supplemental video 4:*

The video shows a *MTFR2* knockout HeLa cell transfected with mCherry-tubulin undergoing mitosis. Note the spindle oscillation and rocking during the prolonged mitosis.

*Supplemental video 5:*

The video shows how control HeLa cells responded to taxol treatment. The movie started 8 hrs after release from thymidine release. Taxol was added at the time of release.

*Supplemental video 6:*

The video shows how *MTFR2* knockout HeLa cells responded to taxol treatment. The movie started 8 hrs after release from thymidine release. Taxol was added at the time of release.

## Acknowledgments

The authors thank Dr. Kam Yeung and Dr. Fan Dong for reagents and discussions on preparing lentiviruses, and Dr. Michael Moenk for developing the lentivirus protocol in our lab. The work was supported, in whole or in part, by National Institutes of Health (R01CA169500 and R15CA238894) and UToledo Biomedical Research Innovation Award to STL.

