## Supplemental figures 1-8 for "MTFR2 regulates mitochondrial fission and impacts spindle integrity during mitosis"

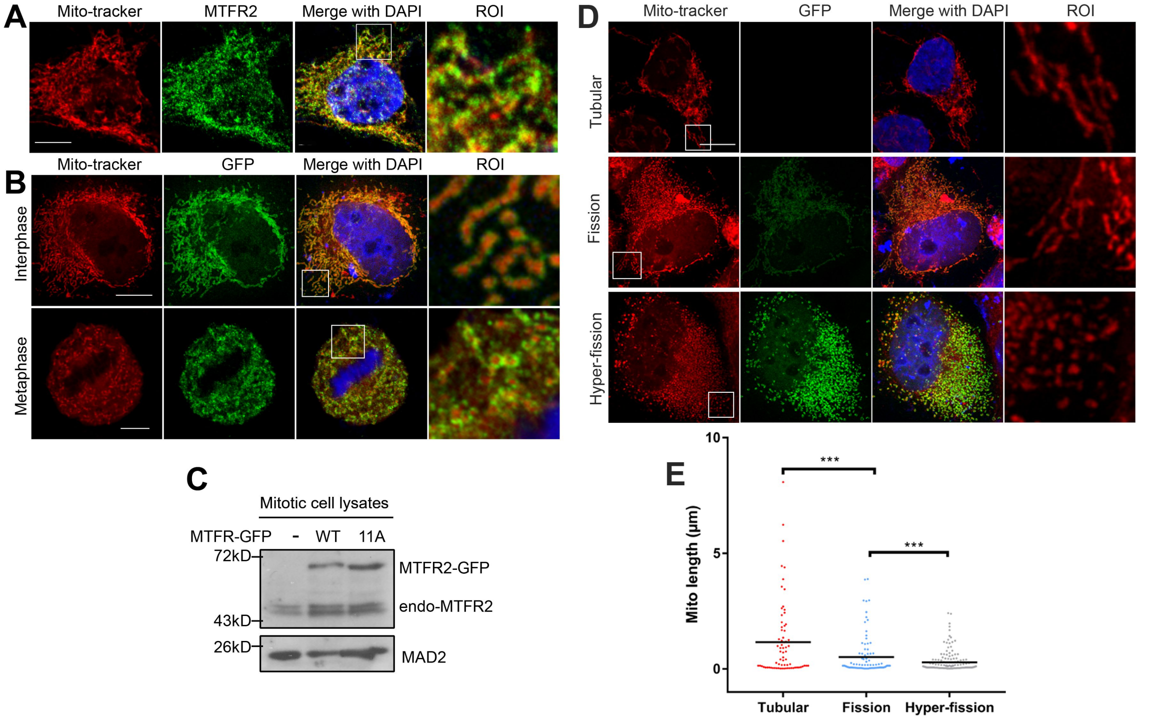

A
Mito-tracker
MTFR2
Merge with DAPI
ROI
B
Mito-tracker
GFP
Merge with DAPI
ROI
Interphase
Metaphase
C
Mitotic cell lysates
MTFR-GFP
-
WT
11A
72kD
43kD
26kD
MTFR2-GFP
endo-MTFR2
MAD2
D
Mito-tracker
GFP
Merge with DAPI
ROI
Tubular
Fission
Hyper-fission
E
Mito length (μm)
10
5
0
***
***
Tubular
Fission
Hyper-fission

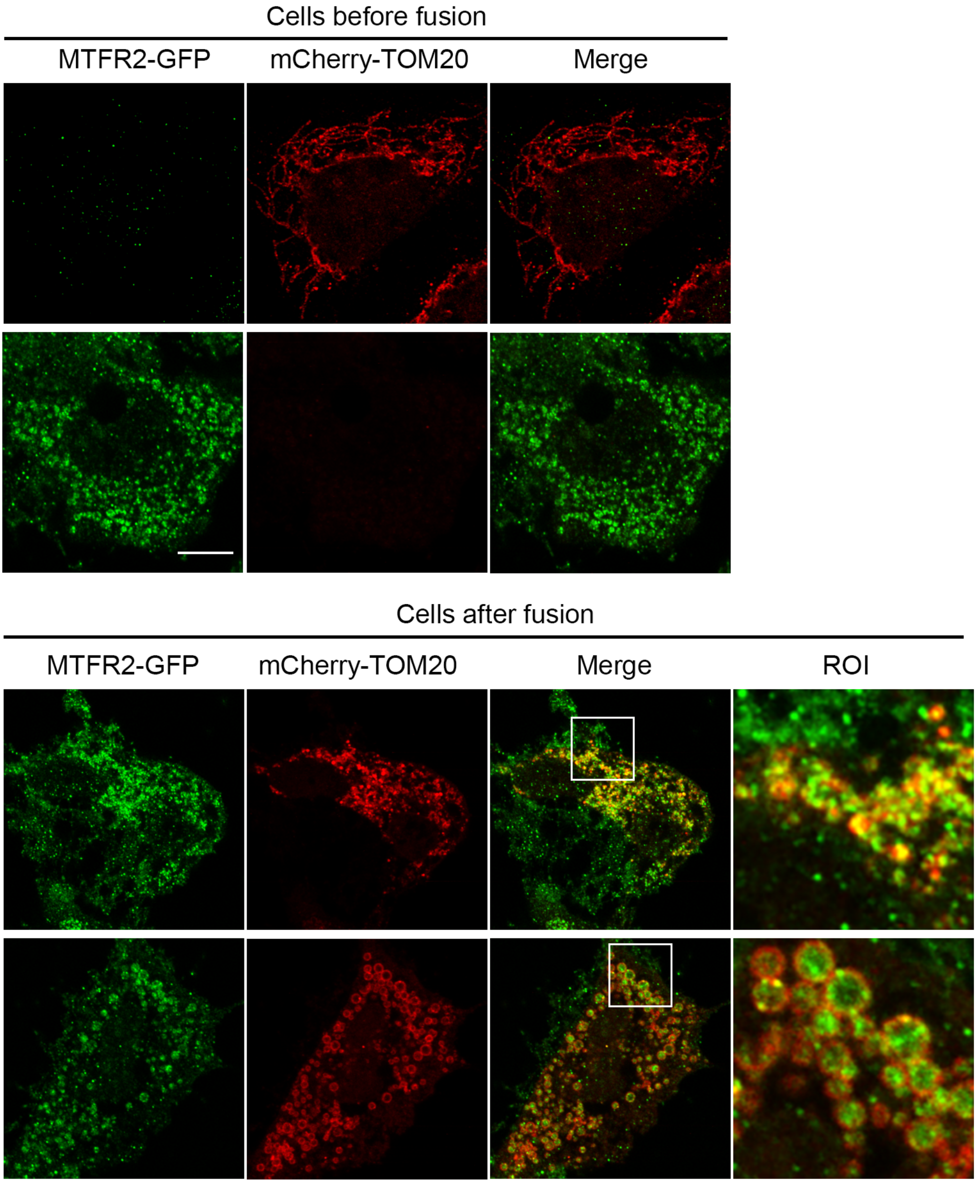

Cells before fusion
MTFR2-GFP
mCherry-TOM20
Merge
Cells after fusion
MTFR2-GFP
mCherry-TOM20
Merge
ROI

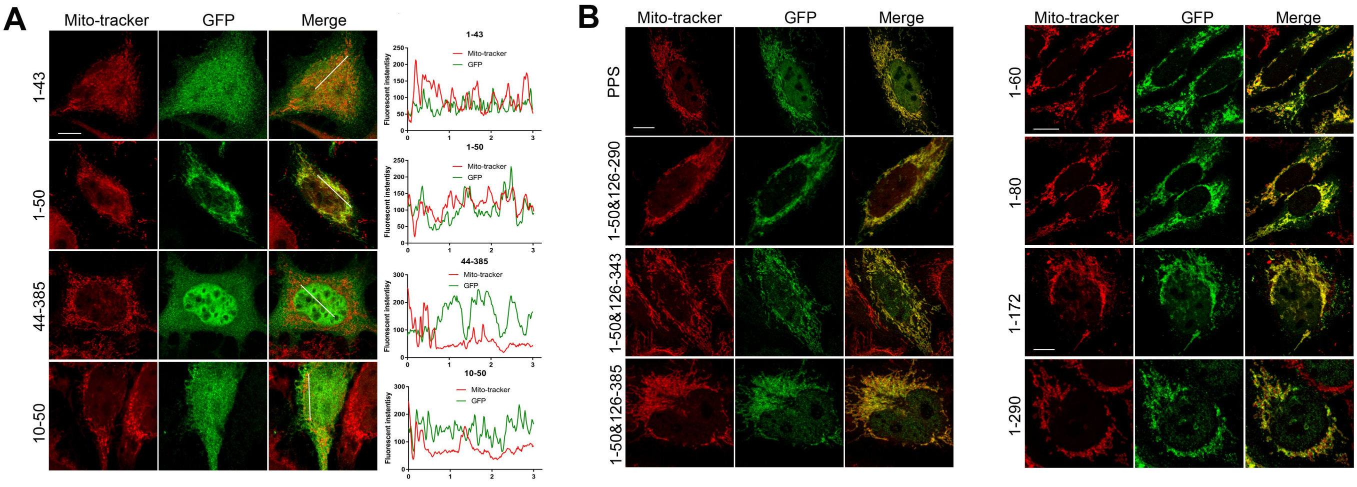


**C**

| MTFR2 Truncations or mutants | Mitochondrial targeting | Mitochondrial fission |
|---|---|---|
| Full length | + | + |
| 1-43 | - | - |
| 1-50 | + | - |
| 1-60 | + | - |
| 1-80 | + | - |
| 1-126 | + | - |
| 1-172 | + | - |
| 1-290 | + | - |
| 10-50 | - | - |
| 44-385 | - | - |
| Q155H | +/- | - |
| Q155E | +/- | - |
| P211R | + | - |
| P211S | + | ->+ |
| R290Q | + | -<+ |
| E126Q | + | -<+ |
| PPS | + | - |
| 1-50&126-290 | + | - |
| 1-50&126-343 | + | - |
| 1-50&126-385 | + | - |

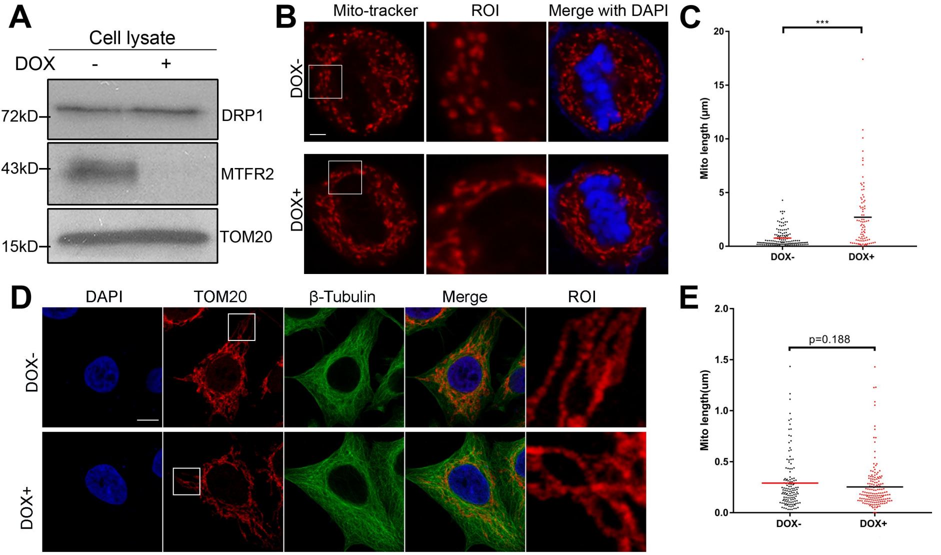
A
Cell lysate
DOX
-
+
72kD
DRP1
43kD
MTFR2
15kD
TOM20
B
Mito-tracker
ROI
Merge with DAPI
DOX-
DOX+
C
***
Mito length (μm)
20
15
10
5
0
DOX-
DOX+
D
DAPI
TOM20
β-Tubulin
Merge
ROI
DOX-
DOX+
E
p=0.188
Mito length(um)
2.0
1.5
1.0
0.5
0.0
DOX-
DOX+

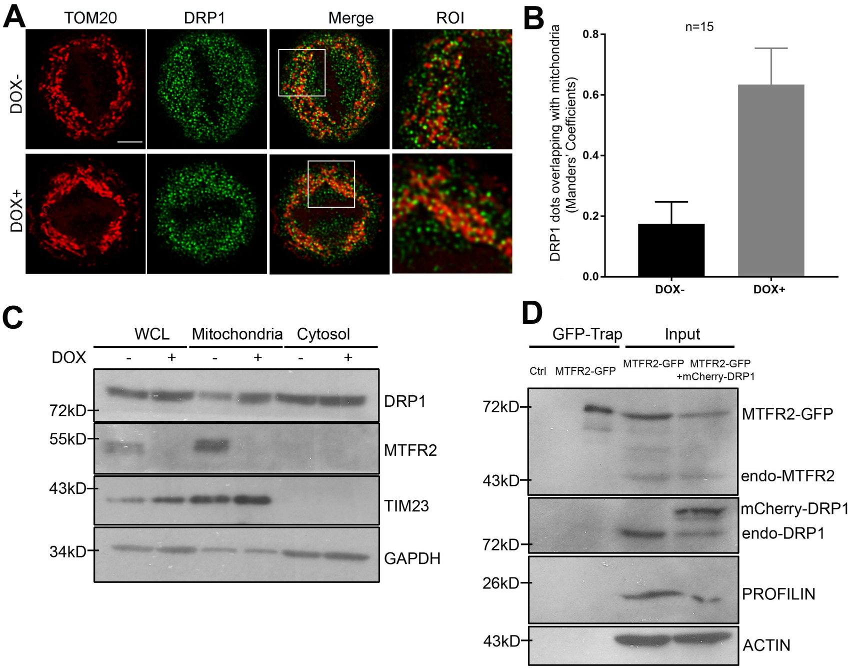
A
TOM20
DRP1
Merge
ROI
DOX-
DOX+
B
n=15
DRP1 dots overlapping with mitchondria
(Manders' Coefficients)
0.8
0.6
0.4
0.2
0.0
DOX-
DOX+
C
WCL
Mitochondria
Cytosol
DOX
-
+
-
+
-
+
72kD
55kD
43kD
34kD
DRP1
MTFR2
TIM23
GAPDH
D
GFP-Trap
Input
Ctrl
MTFR2-GFP
MTFR2-GFP
MTFR2-GFP
+mCherry-DRP1
72kD
43kD
72kD
26kD
43kD
MTFR2-GFP
endo-MTFR2
mCherry-DRP1
endo-DRP1
PROFILIN
ACTIN

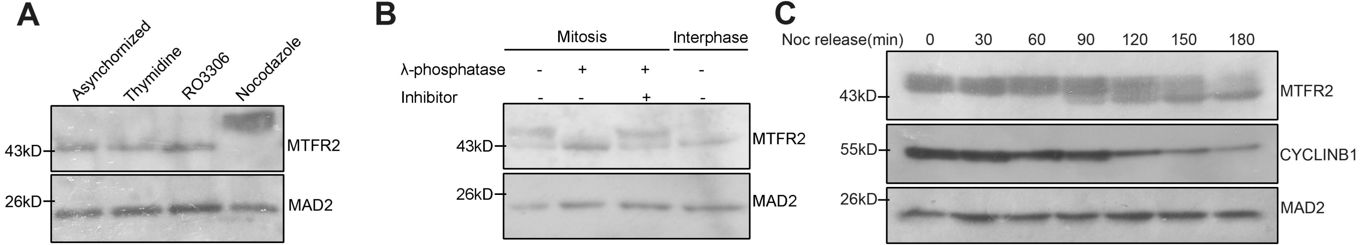
A
Asynchornized
Thymidine
RO3306
Nocodazole
43kD
MTFR2
26kD
MAD2
B
Mitosis
Interphase
λ-phosphatase - + + -
Inhibitor - - + -
43kD
MTFR2
26kD
MAD2
C
Noc release(min) 0 30 60 90 120 150 180
43kD
MTFR2
55kD
CYCLINB1
26kD
MAD2

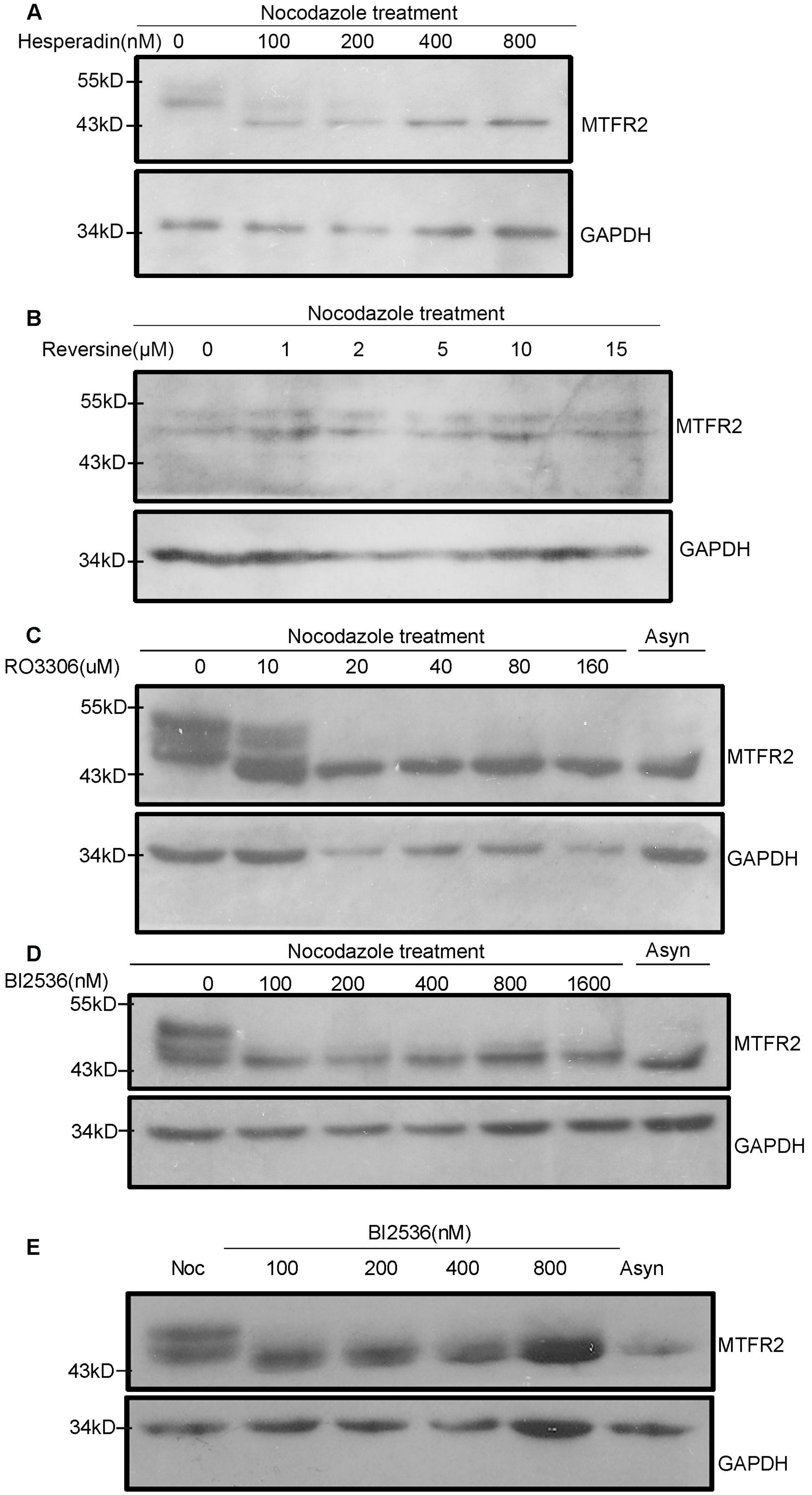
A
Nocodazole treatment
Hesperadin(nM) 0 100 200 400 800
55kD
43kD
MTFR2
34kD
GAPDH
B
Nocodazole treatment
Reversine(μM) 0 1 2 5 10 15
55kD
43kD
MTFR2
34kD
GAPDH
C
Nocodazole treatment
Asyn
RO3306(uM) 0 10 20 40 80 160
55kD
43kD
MTFR2
34kD
GAPDH
D
Nocodazole treatment
Asyn
BI2536(nM) 0 100 200 400 800 1600
55kD
43kD
MTFR2
34kD
GAPDH
E
BI2536(nM)
Noc 100 200 400 800 Asyn
43kD
MTFR2
34kD
GAPDH

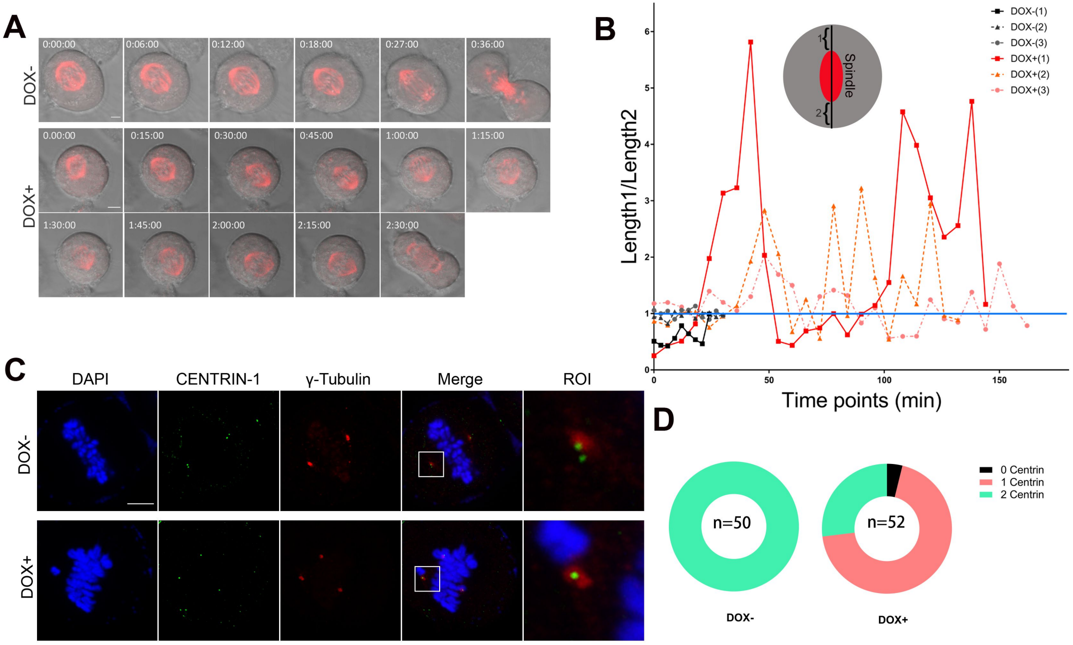
A
DOX-
DOX+
0:00:00
0:06:00
0:12:00
0:18:00
0:27:00
0:36:00
0.00:00
0:15:00
0:30:00
0:45:00
1:00:00
1:15:00
1:30:00
1:45:00
2:00:00
2:15:00
2:30:00
B
Length1/Length2
Time points (min)
Spindle
DOX-(1)
DOX-(2)
DOX-(3)
DOX+(1)
DOX+(2)
DOX+(3)
C
DAPI
CENTRIN-1
γ-Tubulin
Merge
ROI
D
n=50
n=52
DOX-
DOX+
0 Centrin
1 Centrin
2 Centrin
